# Phylogenetic modelling of compositional and exchange rate changes over time

**DOI:** 10.1101/2025.03.14.643246

**Authors:** Peter G. Foster

## Abstract

Changes in the process of evolution occurs over time, including compositional tree heterogeneity (CTH) and exchange rate tree heterogeneity (ERTH). Models that can accommodate CTH and ERTH in molecular evolution are described. Fit of these models was compared using a likelihood ratio test in maximum likelihood, and in Bayesian analysis using the conditional predictive ordinate (CPO)-based log pseudomarginal likelihood (LPML), also leave-one-out crossvalidation (LOO-CV). CTH and ERTH can be flexibly modelled in a Bayesian framework with tree-heterogeneous models that tune themselves to the amount of heterogeneity in the data being analysed.

Since phylogenetic analysis is usually done using tree-homogeneous models, effects of CTH and ERTH on subsequent phylogenetic analysis using such models were described. Compositional effects due to CTH were seen as expected, for example where unrelated taxa with similar compositions would group together in homogeneous analysis. Similar effects were also demonstrated due to ERTH.

Detection of CTH and ERTH by modelling is compared to detection using matched pairs tests (MPTs) that have been used to test molecular sequences for stationarity, reversibility, and homogeneity (SRH). Comparisons between modelling and MPTs on data simulated on very simple trees showed that the two approaches were equivalent, but simulations on larger trees showed that the two approaches differed greatly. Modelling showed greater power, especially in detection of ERTH, and some ERTH was completely invisible to MPTs but was decisively detected by modelling.

Detection and modelling of CTH and ERTH is shown in two empirical examples.

## Introduction

Success of model-based molecular phylogenetic inference depends on whether the models that are used are adequate. While the simplest phylogenetic models had assumptions that were not biologically realistic, phylogenetic models have been improved in various ways, relaxing assumptions to make the models more realistic and better fit the molecular data. This study looks at relaxing the assumption of having an unchanging evolutionary process over time.

A phylogenetic model such as the GTR model has free composition and exchange rate parameters (Tavaré, 1986). These parameters are generally re-optimized for every new phylogenetic analysis. This is necessary in part because these parameters differ in different evolutionary groups, because there are differences in evolutionary process over time. In a similar way there are group-specific empirical amino acid models (Abascal et al., 2007; Adachi & Hasegawa, 1996; Le et al., 2017; Rota-Stabelli et al., 2009; Yang et al., 1998). Again that is because the process of evolution changes over the tree of life. Like the parameters of the GTR model, these empirical amino acid models are composed of a composition component and an exchange rate component; both of these differ between models, and imply that there is heterogeneity in both.

This study describes methods used to detect and models used to accommodate these tree-heterogeneous processes. Detecting by modelling is common in phylogenetics, where molecular sequences are evaluated with and without the model component of interest, and looking for a better model fit when that component is included. That is done here in both a maximum likelihood and Bayesian framework, and models are compared by looking at how well the models fit the data. There are models that accommodate compositional changes over time (Blanquart & Lartillot, 2006, 2008; Foster, 2004; Galtier et al., 1999; Galtier & Gouy, 1998; Yang & Roberts, 1995). Models that accommodate exchange rate changes over time have not been used as much (Foster et al., 2009), and this will be looked at more closely here.

## Methods

The evolution of molecular sequences, DNA and protein, can be described by a continuous-time Markov process. The simplest models, such as the Jukes-Cantor model, can be described with closed-form equations, while more complex models, such as the GTR model, can be described by a rate matrix (Jukes & Cantor, 1969; Tavaré, 1986). As described by Swofford et al. 1996, that rate matrix *Q* can be decomposed into a composition vector *π* and rate matrix *R*. This parameterization is commonly used (Minh et al., 2020; Ronquist et al., 2012; Swofford, 2002), and is used here (Figure 1).

**Figure 1:**
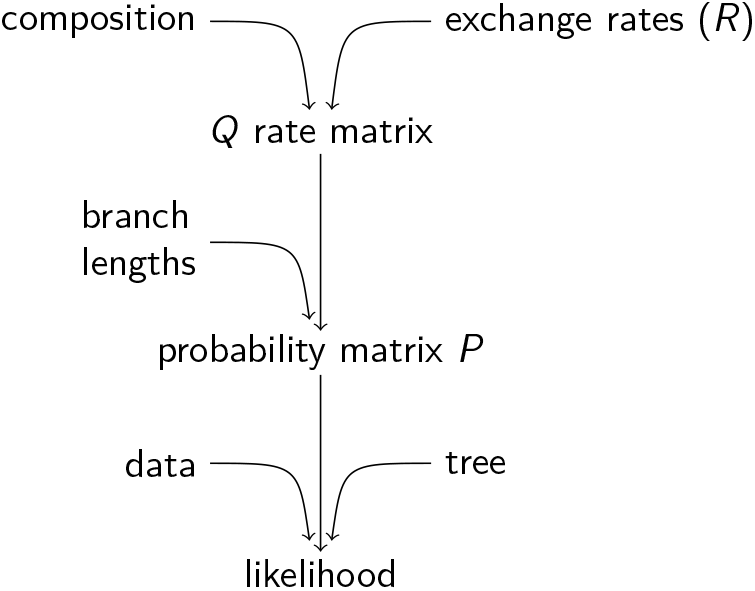
Parameterization context. It is possible for the composition and the *R* exchange rates to differ over the tree. The R-parameters are often described as a matrix, but since the elements on the diagonal are not applicable, and the remaining elements are usually symmetrical, the R-parameters are often described as a short vector. A transition probability matrix *P* can be made from *Q* using the branch length *ν* by *P* = *e*^*Qν*^ . Together with the sequence data and the pruning algorithm on a tree, a likelihood can then be calculated.

Alignments of molecular sequences will often have taxa that have different character state compositions; this is further evidence that the process of evolution changes over time. Here this will be referred to as compositional tree heterogeneity, or CTH (Table 1). This study will also look at changes in exchange rates over time, which will be called exchange rate tree heterogeneity or ERTH. CTH and ERTH can be modelled, and here focus will be on NDCH, NDRH, NDCH2, and NDRH2 (Foster, 2004, 2025; Foster et al., 2009). Changes in the evolutionary process over time can occur anywhere on the tree, gradually or suddenly. This is approximated in the tree-heterogeneous models described here by allowing changes to the model parameters only at nodes, and so “node discrete” in the ND– models described. Between nodes the process is modelled as homogeneous.

**Table 1:** Abbreviations.

|  |  |
| --- | --- |
| CTH | Compositional tree heterogeneity |
| ERTH | Exchange rate tree heterogeneity |
| NDCH | Node discrete compositional heterogeneity model, to model CTH. It has a fixed number of composition vectors over the tree, generally fewer than the number of nodes in the tree; the composition vectors would then be shared over a subset of nodes. As implemented, placement of composition vectors on the tree is fixed in simulations and in ML, but the placement can change in the Bayesian MCMC implementation. Notation such as NDCH(4) would mean having four composition vectors. NDCH-fully means fully parameterized, with a composition vector on all nodes. |
| NDRH | Similar to NDCH, to model ERTH |
| NDXH | NDCH or NDRH |
| NDCH2 | Fully parameterized NDCH, to model CTH, with a prior on the distance from composition vectors to a sampled reference value. |
| NDRH2 | Similar to NDCH2, to model ERTH |
| NDXH2 | NDCH2 or NDRH2 |
| LRT | likelihood ratio test |
| LPML, LOO-CV | Log pseudomarginal likelihood, a measure of model fit used in Bayesian analysis, based on the conditional predictive ordinate (CPO) (Lewis et al., 2014). Also the CPO (cross-predictive ordinate) implementation of leave-one-out cross validation (LOO-CV) (Lartillot, 2023). |
| SRH | An evolutionary model or process that is stationary, reversible, homogeneous. Tree-heterogeneous model components might be locally SRH but not SRH tree-wide. |
| MPT | Matched pairs test |
| MPTS | Matched pairs test for symmetry. Bowker’s test |
| MPTMS | Matched pairs test for marginal symmetry. Stuart’s test |
| MPTIS | Matched pairs test for internal symmetry. Ababneh’s test |

In NDCH and NDRH the models would usually not be fully parameterized over the tree, where each branch or node has a separate set of parameters. Rather, to avoid overparameterization, composition vectors and exchange rate matrices would be shared such that they can be assigned to more than one part of the tree. This is a partially relaxed version of the more usual homogeneous case where the single composition vector and rate matrix are shared by all the nodes over the tree. There are many possible ways that a small number of composition vectors and exchange rate matrices can be assigned on the nodes of a large tree, and as implemented in maximum likelihood that pattern is fixed for each analysis. It would be desirable to allow reassignment and optimization of how the composition vectors and exchange rate matrices are placed on the tree, and the way this problem was addressed in a Bayesian framework was to have MCMC proposals to allow reassignment of different composition vectors or rate matrices to different branches. A problem with both NDCH and NDRH, however, is that the number of composition vectors and rate matrices is fixed at the beginning of an analysis, and usually one would not know how many are needed to adequately model the heterogeneity until running the analysis. It is possible to fully parameterize the tree, with different composition vectors assigned to all the nodes of the tree or have different rate matrices assigned to all branches of the tree. When fully parameterized this way the model should be able to accommodate the maximal heterogeneity allowed by this strategy, although it comes with the possibility of over-parameterization. The problem of over-parameterization is addressed with the NDCH2 and NDRH2 implementations of these tree heterogeneous models, which, in a Bayesian MCMC, use a prior probability on the values of the compositions and exchange rates to decrease the tendency to overfit. In these models the values of the composition vectors or rate matrices are constrained with a Dirichlet prior that tends to keep the proposals more or less close to a sampled reference value. The prior has a hyperparameter that controls the strength of the constraint on the proposals. That hyperparameter would generally be free and sampled, and in this way the model can tune itself to the amount of heterogeneity in the data. In the current implementation these priors are separate between the leaf nodes and the internal nodes.

In a maximum likelihood framework the likelihood ratio test is used to compare models, using the *χ*^2^ approximation to assess significance. A measure of model fit in a Bayesian context based on the conditional predictive ordinate (CPO; cross-predictive ordinate in Lartillot 2023) was described by Lewis et al. 2014, and is used here for model comparison. The CPO is a site-specific measure of model fit that can be combined over sites to make the log pseudomarginal likelihood (LPML), a measure of overall model fit. This method was further described and recommended in a commentary by Lartillot 2023. It is described there as the CPO approach to leave-one-out cross-validation (LOO-CV), and is especially attractive because it is computationally efficient.

Models were implemented in the software p4 (Foster, 2025). DNA models were used in this study, but the software also implements the models for protein and arbitrary (“standard”) datatype.

Methods for matched pairs tests for symmetry, MPTs, were described in Ababneh et al. 2006 The test for symmetry (MPTS, Bowker’s test) can be decomposed to two more specific tests — the matched pairs test for marginal symmetry (MPTMS) and the matched pairs test for internal symmetry (MPTIS). In their usual form these tests are pairwise, testing one pair of sequences, which is how it was done in this study. The methods were re-implemented in p4.

Tree of Life rRNA data was obtained from C.J.Cox (Cox et al., 2008). Phylogenetic analyses of these were done with the data and model partitioned into SSU and LSU partitions. The among-partition rates were fixed ML estimates to facilitate faster convergence.

The alignment from the study of Asian geckos was obtained from Dryad, and the ND2 partition, 1011 sites, was extracted (Brown et al., 2012b, 2012a). The single blank sequence was removed, leaving 40 sequences in the alignment. The remaining sequences ranged from 837 to 1011 nt, with 489 alignment gaps. The Bayesian analysis tree was provided with the data, which was re-rooted on a bifurcating root on the outgroup `Tarentola.mauritanic` for analysis here. The tree had two unresolved internal splits. Bayesisan MCMC analysis was done using the fixed tree, using models GTR, NDCH2, NDRH2, and NDCH2+NDRH2, all with gamma-distributed among-site rate variation (Yang, 1994).

## Results

### Comparing models using LRT and LPML

We assess whether an attribute of evolution is a worthwhile component in a model by comparing models with and without that component. In maximum likelihood we can compare using the LRT, the likelihood ratio test. For simple tree-heterogeneous models, such as the NDCH(2) model with two composition vectors instead of the single composition vector in the GTR model, the null distribution is *χ*^2^-distributed (Supplementary Figure S1).

For larger phylogenetic problems with more complex models it can be better to analyse in a Bayesian framework using an MCMC, and then compare models by comparing LPML values. Using LPML was introduced in a phylogenetic context in Lewis et al. 2014 but has not been commonly used (but see Lartillot 2023). To use it for comparing models to decide whether an increase in LPML is meaningful, we need to know how it behaves for null comparisons such as comparing a well-fitting model with an over-parameterized one. Several such comparisons were made using simulations on four-taxon trees (Figure 2) measuring LPML differences (Figure 3). There it can be seen that for all conditions tested, for most replicates the LPML for the overparameterized MCMC was less than that for the simulation model, showing a penalty for overparameterization. To see if this was also the case with larger trees, some tests were done with 40-taxon trees, under similar conditions as above with GTR simulations and analysis under the simulation model versus overparameterized models (Supplementary Figure S2). The differences were in the range -8 to 4, similar to that shown in the 4-taxon tree results in Figure 3. Although this survey has been limited in scope and may well depend on the context, we can tentatively suggest that an LPML difference of more than about 10 will be considered meaningful.

**Figure 2:**
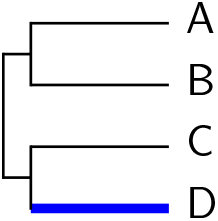
Four-taxon simulation tree. For heterogeneous simulations new model components were placed on the “D” branch.

**Figure 3:**
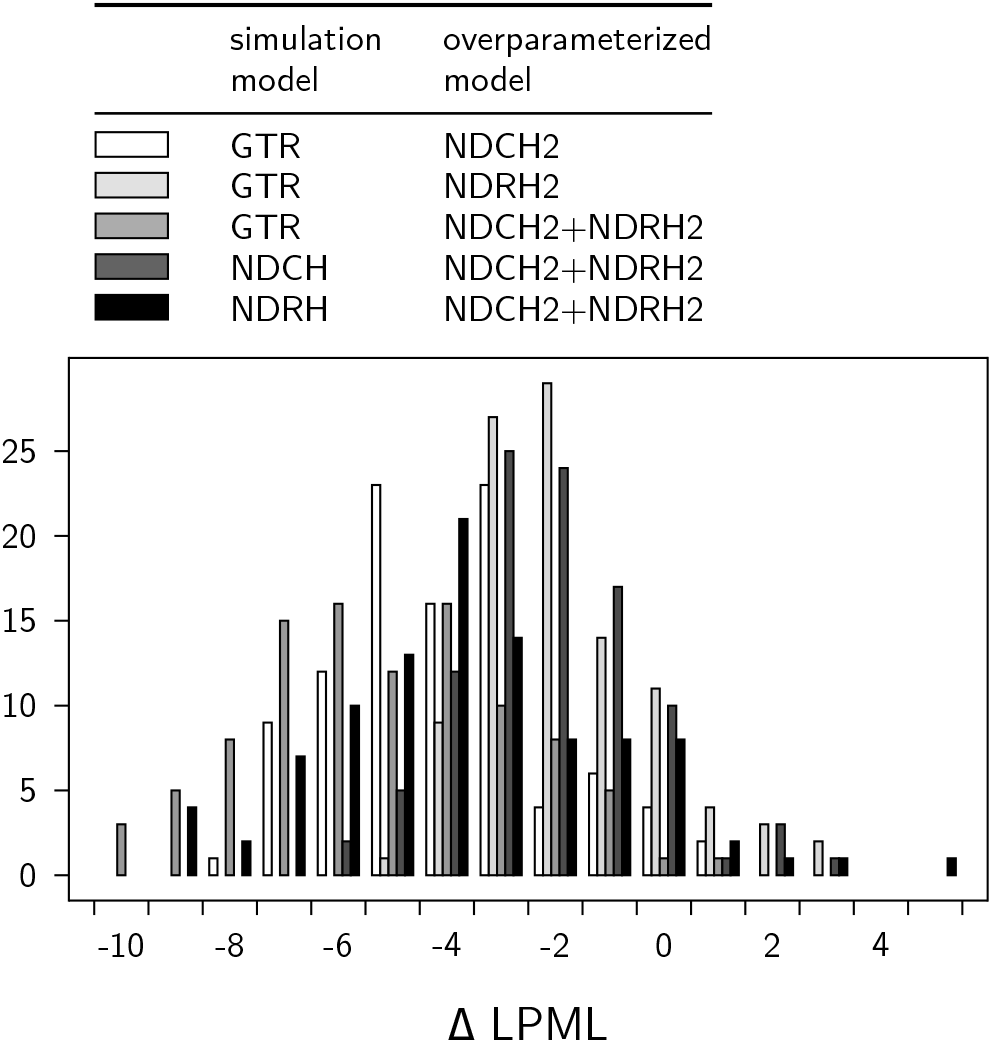
Change in LPML due to overparameterization using data from four-taxon simulations. MCMC analyses were done with the simulation model and with the overparameterized model. The LPML was calculated for both, and the difference plotted (overparameterized - simulation). Each condition used 100 replicates. Most differences showed an LPML penalty for overparameterization.

### Enough parameters without overparameterization

The demonstration here shows the effect of the prior in the NDXH2 (NDCH2 or NDRH2) models, compared to similar models, NDXH-Fully, without the prior. Both the NDXH-Fully and the NDXH2 models are fully parameterized over the tree, meaning that the tree is fully populated with separate composition vectors or exchange rate matrices. Data sets were simulated with small or large amounts of CTH or ERTH and then analysed for model fit using these models (Figure 4). With only a small amount of heterogeneity, either CTH or ERTH, fully parameterized models would be overparameterized, and benefit from the constraint provided by the NDXH2 strategy compared to the NDXH-Fully model (panels A and C). With a large amount of heterogeneity the fit of the NDXH-Fully model is not much worse than the NDXH2 model (panels B and D). Sampled hyperparameters show that the NDXH2 model adjusts itself to the amount of heterogeneity in the data (Figure 5). Larger hyperperameter values exert a stronger constraint on the parameters.

**Figure 4:**
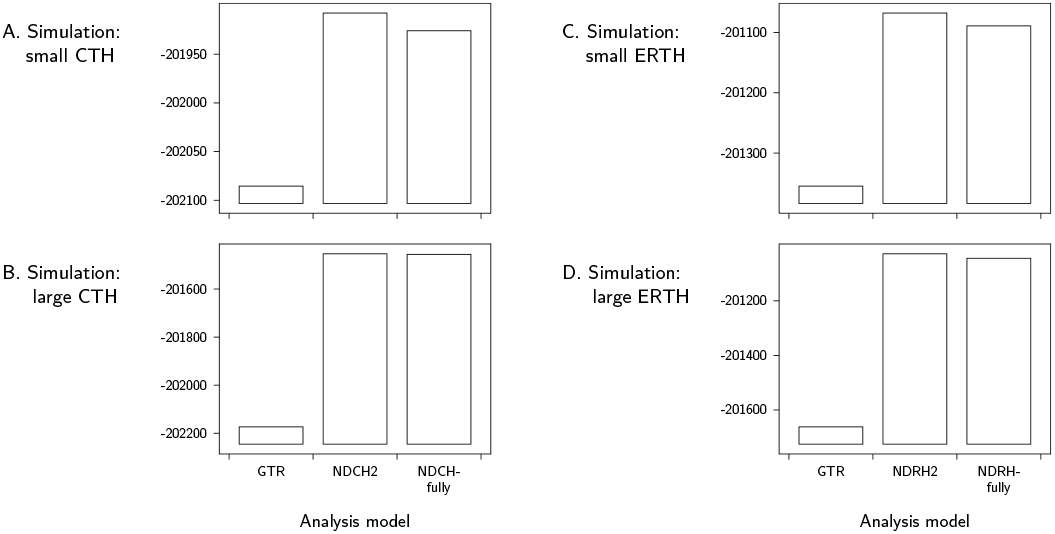
Comparing model fit between NDXH2 and NDXH-Fully. Model fit was measured by LPML. Datasets on an arbitrary 16-taxon tree were simulated containing either a small or a large amount of either CTH or ERTH, and then analysed for model fit on the fixed tree.

**Figure 5:**
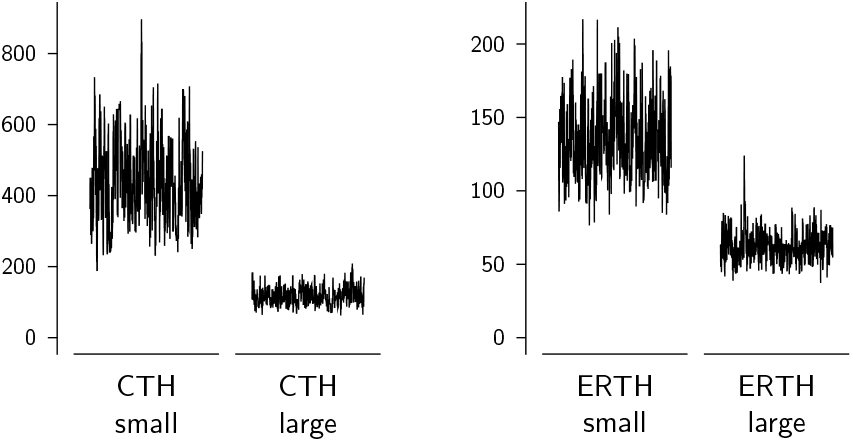
Hyperparameters controlling the constraint in the NDXH2 analyses in Figure 4. Samples of hyperparameters for leaf nodes taken during the MCMC are shown.

### Cross-estimation between CTH and ERTH

The NDCH model was formulated to accommodate CTH only, and the NDRH model was formulated to accommodate ERTH, only. Ideally NDCH would not detect ERTH, nor would NDRH detect CTH; however such cross-estimation is evident. To show this a 4 × 4 matrix of ML analyses was done, using four simulation conditions (with neither CTH nor ERTH, with CTH, with ERTH, and with both) and the corresponding four analysis models. Simulations were done on the four-taxon tree shown in Figure 2. Likelihood ratios were calculated between the GTR analysis and the other three models. Means are shown (Table 2). Likelihood ratios from simulations using the GTR model (row A) have a mean of only a few log units, as expected only showing a slight increase due to overparameterization. Simulations including CTH (row B) show a mean likelihood ratio of 1319 log units when using the NDCH model; such a large value is expected because the data were simulated with CTH. However, those simulations also show a mean likelihood ratio of 96 log units when using the NDRH model, which is unexpected because the NDRH model does not model the CTH in those simulations. Similarly simulations incorporating ERTH (row C) have an expected large mean likelihood ratio of 2742 when using the NDRH model, but also an unexpected mean likelihood ratio of 279 log units when using the NDCH. We can however notice that in the simulations with CTH (row B) the mean likelihood ratio using the NDCH+NDRH model is only 2.5 log units larger than the likelihood ratio from NDCH alone, and does not reflect the addition of the erroneous NDRH contribution. Similarly in the ERTH (row C) simulations, the NDCH+NDRH mean likelihood ratio is only 1.6 log units more than the likelihood ratio from NDRH alone. This observation about the combined NDCH+NDRH model allows us to infer the relative contributions of the NDCH and NDRH when there is cross-estimation as seen here. In the results with simulations with both CTH and ERTH (row D) the mean likelihood ratio using NDCH+NDRH is much greater than either NDCH or NDRH separately, and allows us to infer that these data have both CTH and ERTH.

**Table 2:** Cross-estimation of CTH and ERTH using ML. Four simulations on 4-taxon trees, 2000 replicates each, were evaluated with GTR, NDCH, NDRH, and NDCH + NDRH. Likelihood ratios between the heterogeneous model and GTR were evaluated; means are shown.

| Simulation | NDCH | NDRH | NDCH+NDRH |
| --- | --- | --- | --- |
| A: no CTH, no ERT | 1.54 | 2.47 | 4.03 |
| B: with CTH, no ERT | 1319.36 | 95.99 | 1321.87 |
| C: no CTH, with ERT | 278.70 | 2741.62 | 2743.17 |
| D: with CTH, with ERT | 973.77 | 2425.77 | 3988.15 |

A parallel set of analyses was done in a Bayesian framework using LPML to measure fit of the model (Figure 6). Panel A shows LPML results from analysis of GTR simulations. The LPML for the GTR analysis was the highest. The LPML values for the other three models were, relative to the GTR value, NDCH2: -2.3, NDRH2: -6.0, NDCH2+NDRH2: -9.0 (Supplementary Table S1), which are expected due to overparameterization. Panel B sequences were simulated with CTH, and has a high LPML when modelled by NDCH2. However, there is also an increase in LPML of the NDRH2 model over the GTR model, erroneously showing ERTH. Reciprocally CTH was erroneously detected in panel C by NDCH2 in sequences containing ERTH but no CTH. However, we can notice that in such cases, modelling with both NDCH2 and NDRH2 together appears to behave usefully. In Figure 6 panel B, where some ERTH is erroneously detected, when analyzed with both NDCH2 and NDRH2 together the LPML is only about 6 log units greater than the LPML value for NDCH2 alone. Similarly in Figure 6 panel C, analysis with both NDCH2 and NDRH2 together results in an LPML value that is only about 4 log units greater than the LPML value for NDRH2 alone. This suggests that when there is cross-estimation, by modelling CTH and ERTH separately and together we can estimate both CTH and ERTH, and their relative contributions. Figure 6 D shows an analysis of simulation data that have both CTH and ERTH, where analysis with NDCH2 and NDRH2 together has a model fit greater than either separately, and allows inference of both CTH and ERTH.

**Figure 6:**
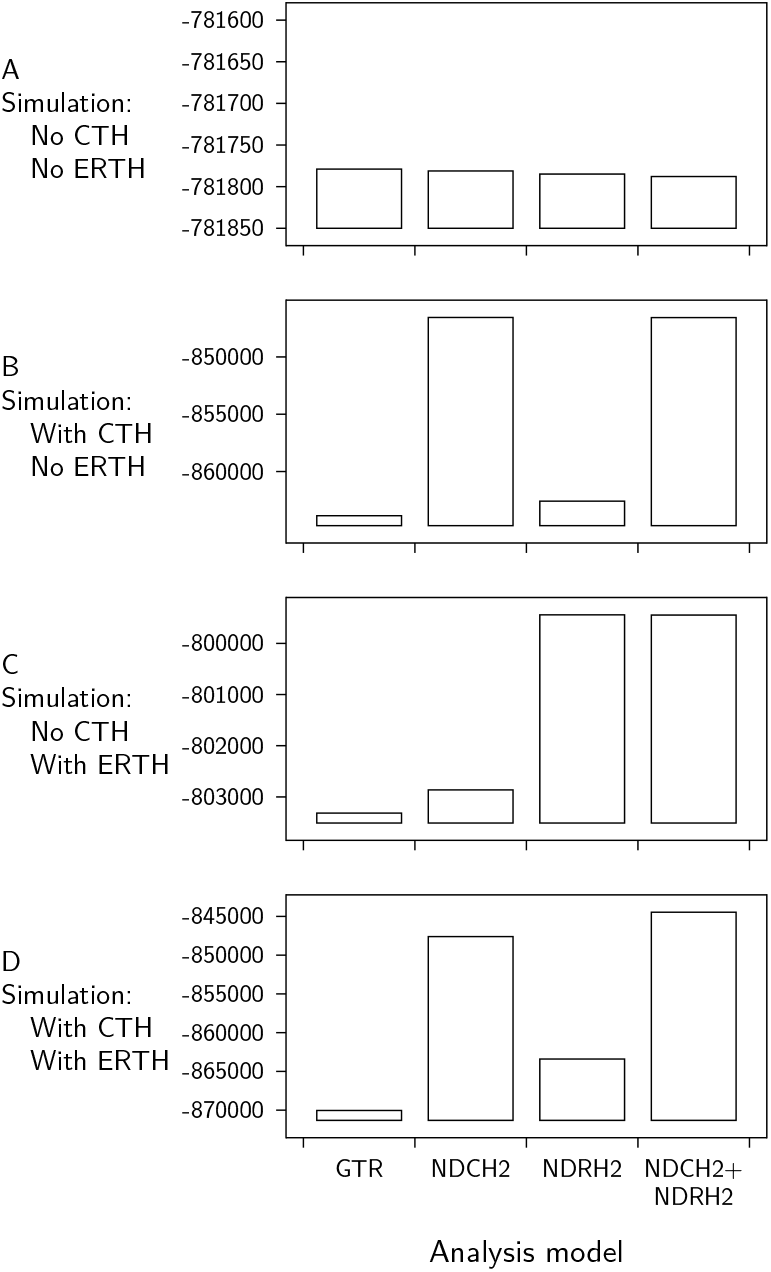
Cross-estimation of CTH and ERTH. Datasets were simulated with and without both CTH and ERTH, and then analysed in an MCMC with and without NDCH2 and NDRH2. Model fit was measured with LPML. Simulations were made on a fixed, arbitrary 7-taxon tree. For simulations incorporating CTH or ERTH, three different fixed composition vectors or three different exchange rate parameter sets were placed on the simulation tree semi-randomly such that each model component was present in at least two locations on the tree.

### Composition and exchange rate effects in simulations analysed with tree-homogeneous models

Usually molecular phylogenetic analysis is done with tree-homogeneous models, and so we would like to know how tree-heterogeneous sequences behave when analysed with tree-homogeneous models. To demonstrate, simulated DNA datasets were generated on a tree with CTH or ERTH in various patterns over the simulation tree (Figure 7, left column). Results in the middle column of that Figure show tree-homogeneous analysis, showing topological distortions.

**Figure 7:**
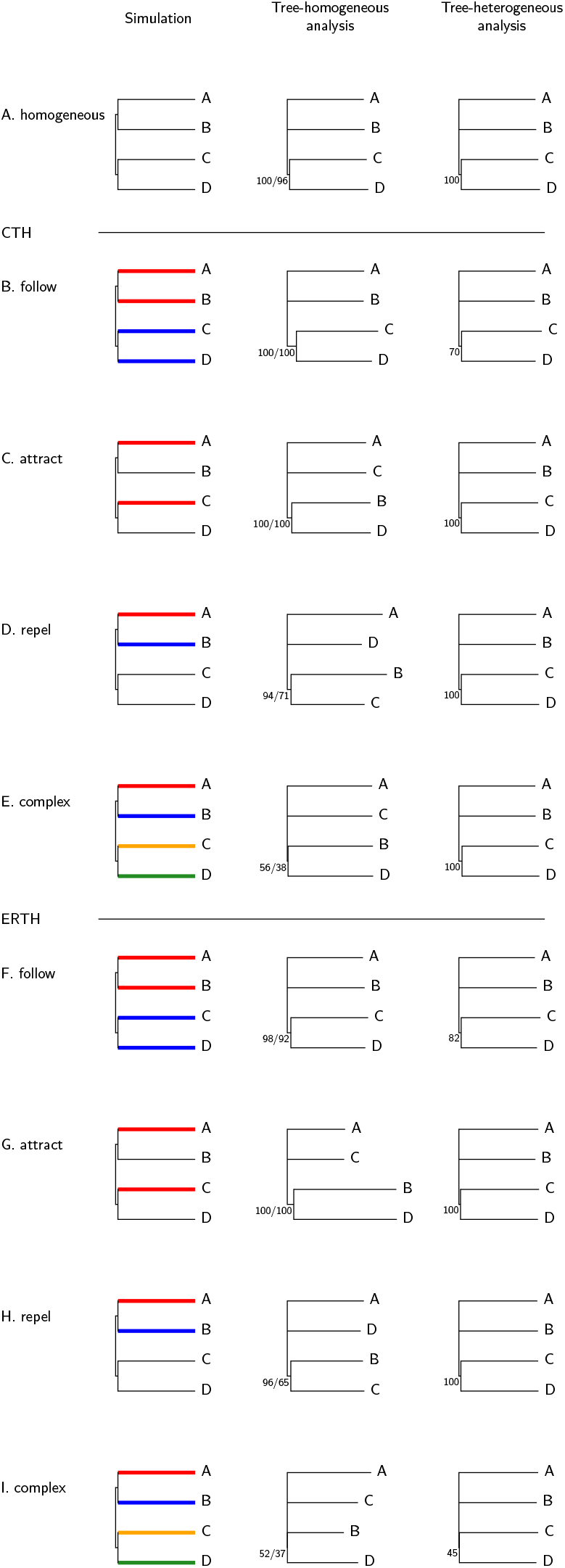
Effects of composition and exchange rate differences over the tree on subsequent analysis of simulated DNA datasets. Analysis used a tree-homogeneous model (middle column), or a treeheterogeneous model (right column). Coloured branches on the simulation trees show the CTH (Rows B – E) or ERTH (Rows F – I) that was superimposed on the background Jukes-Cantor evolution shown in black. Resulting datasets were then analysed with a GTR model using either an MCMC in p4 or maximum likelihood and bootstrap with IQTree with results shown in the homogeneous analysis column (support shown as posterior probability percent/bootstrap percent). The datasets were also analysed using tree heterogeneous models with results as shown in the column on the right; the model was NDCH2+NDRH2 in all cases.

It may be that the most common and expected effect occurs when CTH or ERTH differences *follow*, from the root to the leaves, the same evolutionary path as the tree (Figure 7 rows B, F). In this case we have attraction of sister taxa, which tends to raise support for grouping the sisters in homogeneous analysis. Of the effects shown, this one might be considered the most benign because it increases support for groupings that should be together, but it still must be considered a distortion because it increases support undeservedly. Likely the most well-known effect shown is *compositional attraction*, where taxa that are remote on the tree but with similar sequence compositions are attracted in analysis (Figure 7 row C). Such an attraction can also occur due to ERTH, without any CTH (Figure 7 row G).

When sister taxa differ in composition or exchange rates, they can *repel* each other (Figure 7 rows D, H). Such repulsion can be considered as non-specific attraction to some other part of the tree. The *complex* scenario described in Figure 7 rows E and I is meant to describe an interplay of attractions and repulsions. While it would be easy to predict the distortions from the *follow* or *attract* scenarios on homogeneous analysis, it would be hard to predict the analysis topology from a *repel* or *complex* pattern.

The simulated alignments shown were also analysed with an NDCH2+NDRH2 model, which recovered the simulation topology in all cases (right column in Figure 7) together with a better model fit (Supplementary Table S2). For those analyses involving CTH, posterior predictive simulations show that the model measured with the heterogeneous analysis fit the composition while the homogeneous analysis did not (Supplementary Table S3).

Looking more closely at Figure 7 row B, we can see that the model fit is much better in the tree-heterogeneous analysis compared to the tree-homogeneous analysis (LPML increased by 1094.8, Supplementary Table S2, Row B). This is an example of an analysis that had decreased support for topological resolution (from 100% to 70%), yet was a better fit.

### Cross-estimation is not seen with matched pairs tests as it is with models

Matched pairs tests (MPTs) have been described and advocated as a way to test SRH — stationarity, reversibility, and homogeneity — in molecular sequences (Ababneh et al., 2006). The MPTs are a family of three related tests. The most general MPT is Bowker’s test for symmetry, but it can be decomposed into two more specific tests — the test for marginal symmetry (MPTMS) that tests for stationarity, and the test for internal symmetry (MPTIS). Here simulated alignments are tested using the MPTMS and MPTIS, and compared to testing using modelling and the LRT as done above. These strategies, the LRT and MPTs, have similar goals, and although the methods differ greatly we would expect them to agree.

MPTs are performed on aligned sequences; they do not need trees, models, or a phylogenetic analysis. This means that they can be used before phylogenetic analysis — an advantage over model comparison to identify CTH and ERTH. This has been incorporated into a suggested phylogenetic protocol that involves screening for alignments or sequences that do not meet the assumptions of the proposed analysis methods (Jermiin et al., 2020). However, in their usual form the MPTs are performed on pairs of sequences in an alignment, while what we generally want is an assessment on an entire alignment. Using MPTs on many pairs in an alignment potentially leads to problems with multiple comparisons (Ababneh et al., 2006). Here the MPTs are applied to simulated data where heterogeneity of the pairs is known in advance, and only a single pair of sequences is measured with the MPTs, thereby avoiding multiple comparison.

We first ask whether the MPTs detect CTH and ERTH, and whether they suffer from the problem of cross-estimation of CTH and ERTH as described above for model-based comparison. To do this the MPTMS and MPTIS were applied to the simulations used in Table 2. Mean P-values for those tests are shown in Table 3, and are either zero or about 0.5. P-values close to 0.50 are because they form a uniform distribution, with no significance except for the Type-I error rate. Mean P-values of 0.0 reflect significance — the MPTMS is detecting CTH and the MPTIS is detecting ERTH. If there was cross-estimation then then we would expect to see lowering of the mean P-value from the MPTIS of the B-series simulations, and we would expect to see lowering of the mean P-value from the MPTMS of the C-series simulations. However, there is no evidence in Table 3 for cross-estimation as seen above with with models (Table 2, Figure 6).

**Table 3:** Datasets simulated under conditions of CTH and ERTH were tested with MPTs. Mean P-values over 2000 replicates are shown.

| Simulations | P MPTMS | P MPTIS |
| --- | --- | --- |
| A: no CTH, no ERT | 0.49 | 0.50 |
| B: with CTH, no ERT | 0.00 | 0.52 |
| C: no CTH, with ERT | 0.50 | 0.00 |
| D: with CTH, with ERT | 0.00 | 0.00 |

### Power of MPTs compared to LRT

Statistical power of the MPTMS was first compared to the LRT using simulations on a one-branch, two-taxon tree. To challenge the sensitivity of these tests, simulations were made with a small amount of CTH, such that the mean P-value over 2000 replicates was 0.29 for both tests, and then the two tests were compared using PP-plots (Figure 8, Panel A). The P-values of the two tests were almost identical, differing by less than 10^*−*5^ each on average, showing that the two tests had the same sensitivity to detect CTH under these conditions. Panel B uses the same data points as in panel A, but shows the P-value of each MPTMS plotted against the P-value of the LRT for the same simulation, again showing complete agreement.

**Figure 8:**
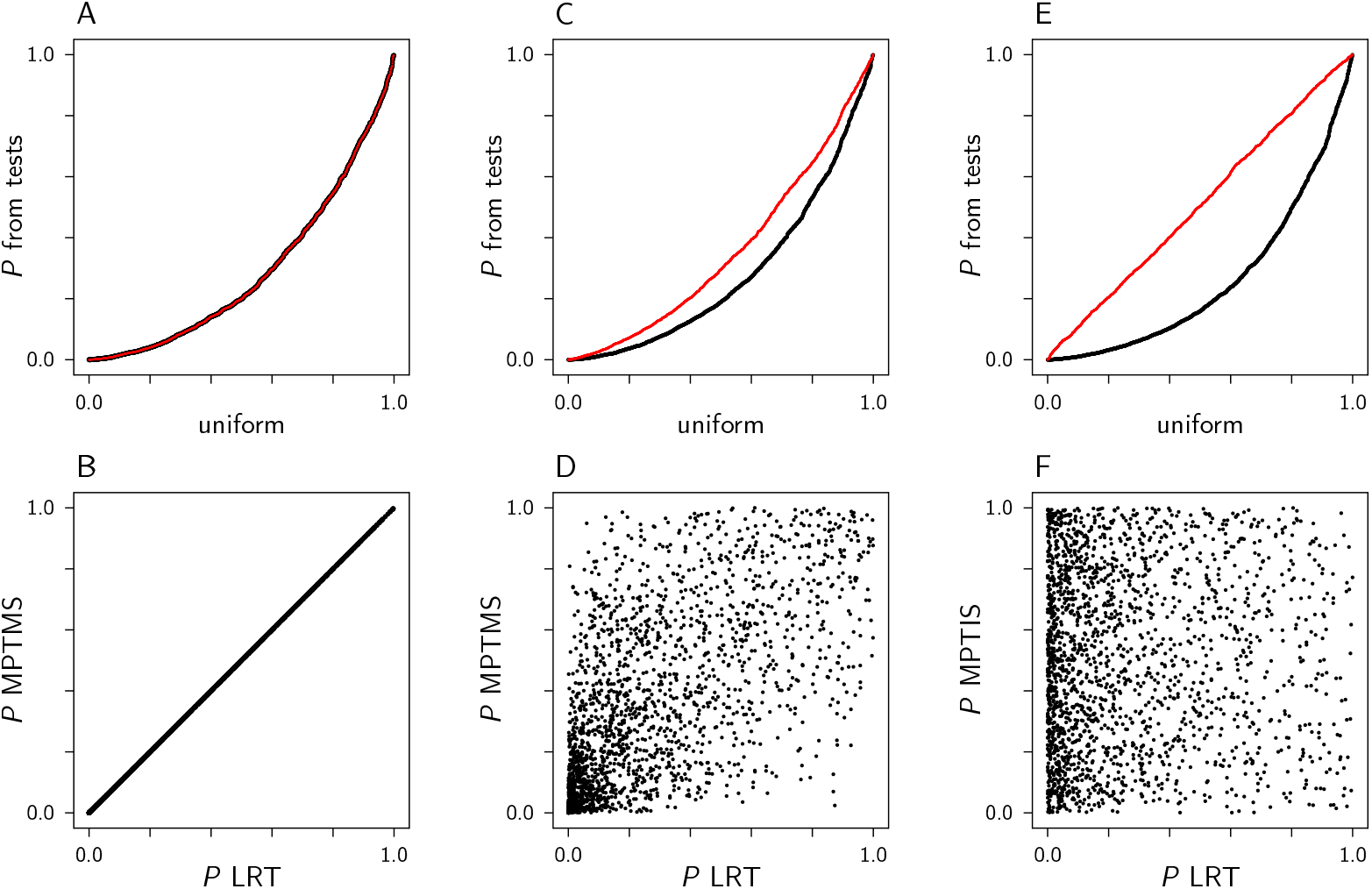
Comparing the statistical power of the MPTs with the LRT. Two thousand replicate alignments, each of length 100000, were made for each condition. Panel A shows a PP-plot from simulations on a one-branch, two-taxon tree with a small amount of CTH, and shows the LRT P-values in black and the MPTMS P-values in red, both plotted against a uniform distribution. Panel B uses the same data points as in panel A, but plotted simulation-by-simulation. Panels C–F are from simulations on a four-taxon tree, with a small amount of CTH in panels C and D, and a small amount of ERTH in panels E and F. Panel C is a PP-plot showing LRT P-values in black and MPTMS P-values in red, both plotted against a uniform distribution. Panel D uses the same data points as panel C, but plotted simulation-by-simulation, showing that evaluations of the two tests for each simulated alignment differed greatly. Panels E and F are similar to panels C and D, but using an LRT using NDRH(2) and the MPTIS to detect ERTH.

Then the power of the MPTs were compared to that of the LRT using simulations on four-taxon trees as in Figure 2, with results shown in Figure 8 panels C–F. To test the power to detect CTH, simulations were made with a small amount of CTH placed on leaf D in the simulation tree. Then the MPTMS was compared to the LRT using GTR versus NDCH(2) in a PP-plot, showing that the LRT was more sensitive (Figure 8 Panel C). When the comparison was made dataset-by-dataset it is seen that the two tests greatly disagreed (Panel D). Although comparing the tests this way from simulations on a one-branch tree showed compete agreement (panel B), this was not so when done with simulations on a four-taxon tree (panel D). Panel E simulations similarly contained a small amount of ERTH on the branch leading to taxon D in the simulation tree, and compares the MPTIS with an LRT of GTR versus NDRH(2) using a PP-plot, showing that the LRT was in general much more sensitive than the MPTIS here. Again the point-by-point individual analyses greatly disagreed with each other (Panel F).

### Some ERTH is invisible to MPTIS

In comparing the MPTs with the LRT it became evident that some kinds of simulated data that contain significant amounts of ERTH, as simulated by the NDRH model and as assessed with the LRT, are such that the ERTH is not visible to the MPTIS (Figures 9 and 10). In the first example, one pair of R-matrices shows ERTH with both the LRT and the MPTIS, but a similar pair shows ERTH with the LRT but not with the MPTIS.

**Figure 9:**
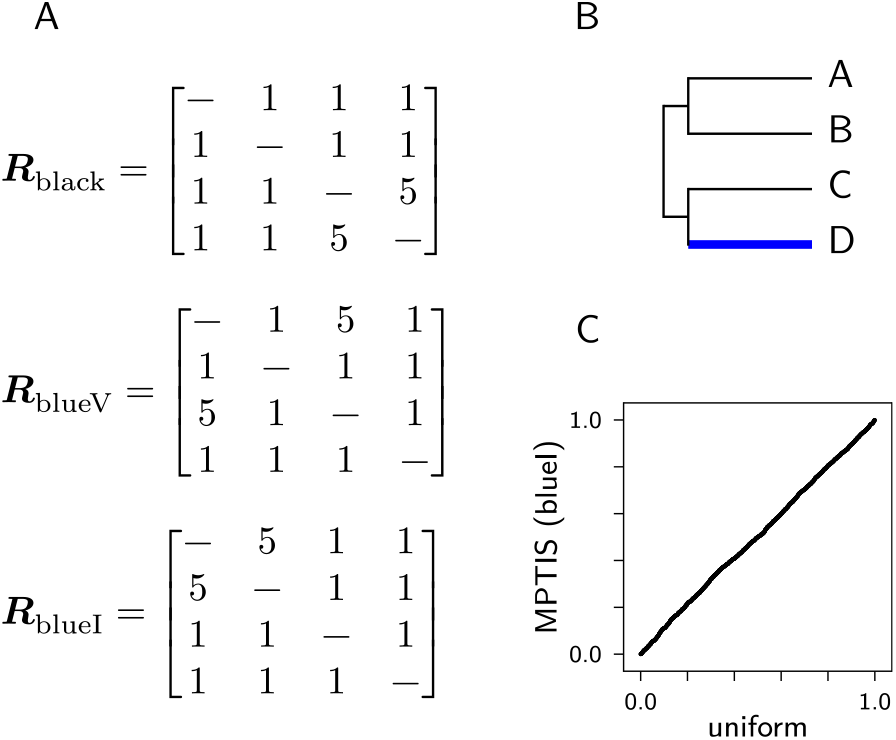
Some ERTH is invisible to the MPTIS. The *R*-matrices used were as shown in Panel A, where the order of the nucleotides is A, C, G, T. For the first test they were placed on the tree such that *R*_blueV_ (*V* for visible) was on the blue branch and the *R*_black_ was on all the other branches (Panel B). Composition was 25% for each character state. Under these simulation conditions datasets are made with ERTH but no CTH, and the resulting ERTH was significant with both the LRT and using the MPTIS (P=0 for all 2000 replicate simulations for both tests). However, if the *R*_blueV_ was replaced with the *R*_blueI_-matrix (*I* for invisible), simulated datasets were made that remain significant with the LRT (P=0 for all 2000 replicate simulations) but with the MPTIS test were significant only at the Type-I error rate, as shown in the PP-plot in Panel C.

**Figure 10:**
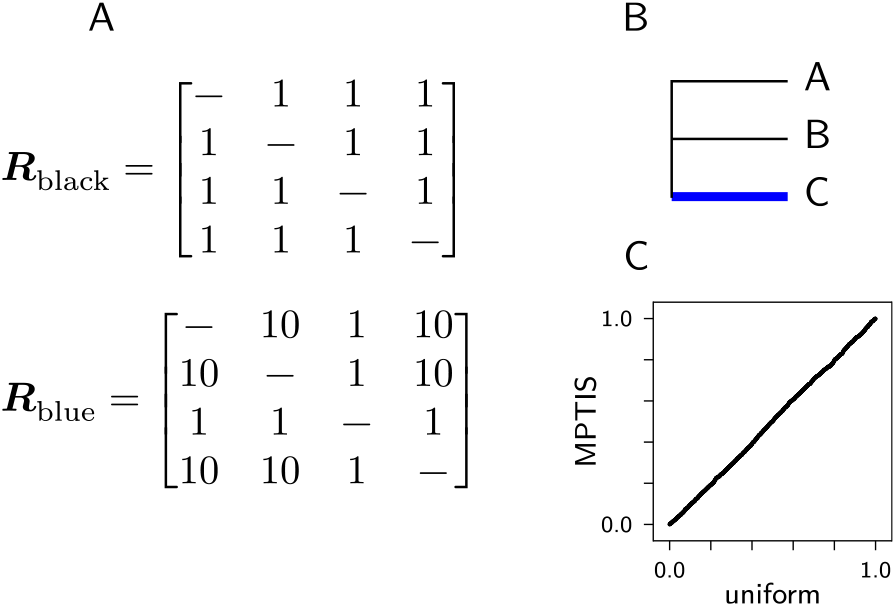
Another example of ERTH that is invisible to the MPTIS. Simulations were made on the three-taxon tree shown in Panel B with R-matrices as shown in Panel A (with nucleotide order A, C, G, T) placed on the tree such that *R*_blue_ was placed on the blue branch and *R*_black_ was placed on the other two branches. Compositions were equal (25% each character state) and there was no CTH. LRTs were done with GTR versus NDRH, and all 2000 replicates were significant (P=0 for all). However, the MPTIS P-values were uniform as shown in the PP-plot in Panel C.

### Example using Tree of Life rRNA

This example uses an rRNA dataset composed of concatenated SSU and LSU genes spanning the Tree of Life (Cox et al., 2008). This study revived the *eocyte hypothesis* for the origin of eukaryotes (Rivera & Lake, 1992). It was analysed in Cox et al. 2008 with the GTR and the NDCH model, and it is re-analyzed here.

This dataset was analysed in Cox et al. 2008 with the GTR model to show a “three-domains” tree of life with 73% posterior probability (as in Figure 11 A, which has that tree with 68% support). They then used the NDCH(2) model, which showed 75% support for the grouping eocytes with eukaryotes. The repeat here using the NDCH2 model showed 96% support for that split (Figure 11 B). Figure 12 shows a higher LPML for NDCH2+NDRH2 than for either NDCH2 or NDRH2 alone, meaning that the data have both CTH and ERTH (The difference between the LPML for NDCH2 and NDCH2+NDRH2 is 63.1 log units; Supplementary Table S4). Using NDRH2 alone shows 60% support for eocytes plus eukaryotes, and using NDCH2+NDRH2 increases support for that grouping to 99%, higher than either NDCH2 or NDRH2 separately. It appears that both CTH and ERTH contribute to the GTR model resulting in the topology shown in Figure 11 A. As measured by posterior predictive simulation using the *X*^2^ test quantity, model fit of the composition of the models used for the rRNA analyses showed that the models using NDCH2 fit the data, while the GTR and the NDRH2-only models did not (Supplementary Table S5).

**Figure 11:**
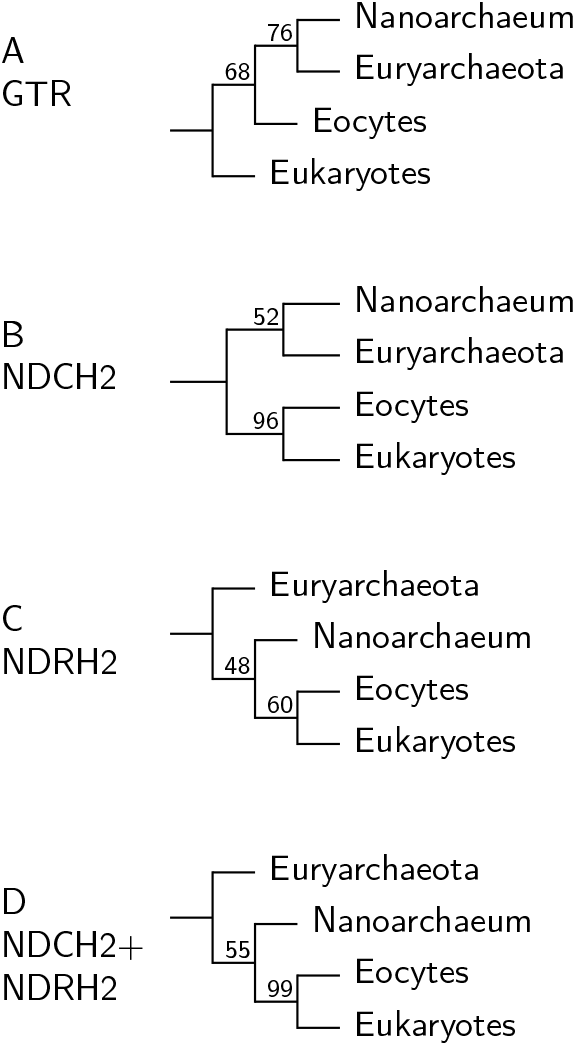
Topologies obtained for different models used in Cox et al (2008) rRNA re-analysis. Trees are rooted on Bacteria.

**Figure 12:**
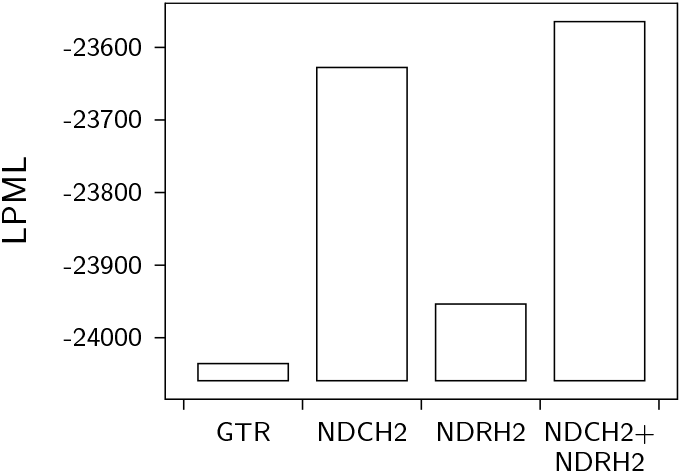
Analysis of Cox et al rRNA. This shows evidence for both CTH and ERTH.

### Example using Asian gecko ND2

This reanalysis is from a study by Brown et al. 2012b, 2012a. While the Tree of Life example above used highly diverged sequences and might therefore be expected to show CTH and ERTH, the study in this example used the ND2 gene from a few genera of asian geckos, and so are from a much smaller taxonomic group than the Tree of Life rRNA example above. Pearson’s *χ*^2^ test rejected compositional homogeneity (P=0) for this dataset. This dataset, including two more short genes, was previously examined using the MaxSymTest as implemented in IQTree, finding some compositional heterogeneity by the MPTMS, but not failing the MPTIS (See Figure 2 in Naser-Khdour et al., 2019). LPML values to show model fit are shown in Figure 13, and show that the LPML using NDCH2+NDRH2 is greater than the LPML for NDCH2 alone by 23.3 log units (Supplemental Table S6), meaning that the dataset has both CTH and some ERTH. Posterior predictive simulations using the *X*^2^ test quantity to measure model fit showed that the models that included NDCH2 fit the composition of the data, but the GTR model and the NDRH2-only model did not (Supplemental Table S7).

**Figure 13:**
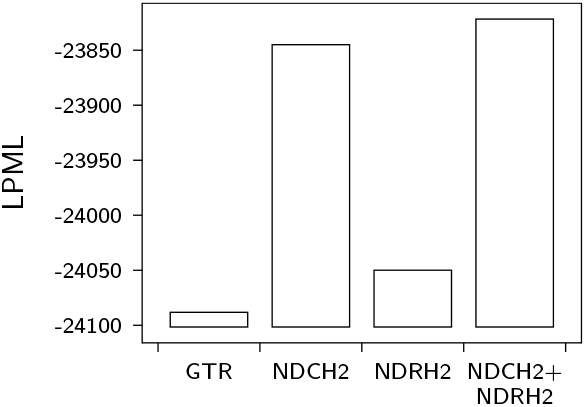
Gecko ND2 analysis.

## Discussion

This study demonstrates phylogenetic modelling of both compositional and exchange rate treeheterogeneity (CTH and ERTH), using tree-heterogeneous NDCH and NDRH models both for simulation and for analysis under maximum likelihood, and NDCH2 and NDRH2 for use in Bayesian analyses. The NDXH2 models, NDCH2 and NDRH2, are fully parameterized over the tree, but avoid overparameterization by constraining the CTH and ERTH parameters, and so have a better fit to tree-heterogeneous data than a similar model without that constraint (Figure 4, NDXH2 versus NDXH-Fully)

Models were compared in maximum likelihood using the likelihood ratio test. In a Bayesian analysis models were compared with the log pseudomarginal likelihood, LPML (also LOO-CV). The LPML was introduced to phylogenetics in Lewis et al. 2014, but has has not been commonly used. Tests to become more familiar with its behaviour show that it often shows a penalty for overparameterization (Figure 3). Posterior-predictive simulations were useful in assessment of model fit, and were used in this study (for example Supplementary Table S3). They have the advantage of providing an assessment of absolute fit of the model to the data (Bollback, 2002; Foster, 2004). However, here these were only used to assess fit of models to CTH because a test quantity to measure compositional heterogeneity, *X*^2^, was at hand. Without a test quantity for ERTH, posterior predictive simulations for ERTH were not done here.

Cross-estimation of CTH and ERTH was seen using the tree-heterogeneous models. This was seen both with the LRT in ML, and with LPML in Bayesian analysis (Table 2, Figure 6).

However, by modelling CTH and ERTH both separately and together it is possible to infer the relative contributions of CTH and ERTH. This cross-estimation is not seen when using MPTs (Table 3).

Models in common use are tree-homogeneous. When used with these models data that have CTH and ERTH may lead to poor parameter estimates and topological distortions (Figure 7). For example, compositional attraction was shown, where unrelated taxa with similar compositions are attracted to one another when using homogeneous analysis. A parallel effect with ERTH was also shown, where taxa with similar exchange rates are attracted to each other, showing a previously unknown way for phylogenetic analysis to go wrong when mis-modelled. Another pattern seen was erosion of support as the model fit improved, when the poorly-fitting tree-homogeneous model had erroneously high support, and the better-fitting heterogeneous model decreased support (Figure 7 Rows B and F). This is a reminder that the goal is accurate estimation, not simply high support for topological resolution, and that a model that fits well benefits parameter estimation as well as topology estimation.

Detection of CTH and ERTH was compared between MPTs and model-based LRTs. It is shown that the MPTMS will detect CTH, and the MPTIS will detect ERTH (Table 3). An advantage of MPTs is that they can be used before phylogenetic analysis, and this would allow identification and removal of problematic taxa or alignments (Jermiin et al., 2020). However, since CTH and ERTH are features of evolution, focussing only on tests that flag violations of SRH with the intention of subsequent exclusive use of tree-homogeneous models is methodologically incomplete, because we may want to be able to also assess fit of tree-heterogeneous models to tree-heterogeneous data. Model-based comparisons are alignment-wide, which is generally what is wanted, while the MPTs are done between sequence pairs, complicating alignment-wide screening (see for example the solution described in Naser-Khdour et al. 2019). As mentioned above, MPTs do not suffer from cross-estimation of CTH and ERTH as do model-based comparisons (Table 2 compared to Table 3), which makes interpretation of the MPTs more direct.

It would be expected that MPTs and model-based detection of CTH and ERTH would agree, and this is true in comparisons testing CTH on simulations on one-branch, two-taxon trees (Figure 8, panels A and B). However, the two approaches differ when tested on four-taxon trees (Figure 8 panels C – F). In the four-taxon tests the LRT was found to have more statistical power in general. Furthermore, comparing these approaches simulation-by-simulation show that MPTs and modelling are very different (Figure 8 D,F). The two approaches differ to the extent that some simulations that contain significant ERTH as measured by the model-based LRT are invisible to the MPTIS (Figures 9, 10).

ERTH was seen in the Tree of Life rRNA dataset, which might be expected because those taxa were highly diverged (Figure 12 panel A). However, we also measured some ERTH, as well as substantial CTH, in the gecko alignment, the taxa of which are more closely related (Figure 13). ERTH was found to be extensive in a survey made using a version of the MPTIS (Naser-Khdour et al., 2019). Modelling found some evidence for ERTH in gecko ND2, while it was not found by Naser-Khdour et al. 2019, possibly because modelling is more sensitive (Figure 8 panel E), or different in other ways (Figure 8, panel F).

There is compelling potential for further improvement of phylogenetic models when we note that biological sequences will generally have compositional heterogeneity over alignment sites (Lartillot & Philippe, 2004). We can model that with profile-based models, such as the CAT model (Lartillot & Philippe, 2004). It would be useful to accommodate CTH and ERTH as well as among-site compositional heterogeneity in the same model (Blanquart & Lartillot, 2008; Feuda et al., 2017; Naser-Khdour et al., 2019).

## Notes

### Competing Interest Statement

The authors have declared no competing interest.

