## Supplementary Information for "Phylogenetic modelling of compositional and exchange rate changes over time"

Supplementary material

#### Contents


Null distribution of the likelihood ratio is $\chi^2$-distributed 2

LPML effects due to overparameterization using 40-taxon tree simulations 3

Cross-estimation between CTH and ERTH 4

Composition and exchange rate effects in simulations analysed with tree-homogeneous models: LPML 4

Composition and exchange rate effects in simulations analysed with tree-homogeneous models: posterior predictive simulations 5

Cox et al RRNA 6

Brown et al 2012 ND2 analysis 7

### Null distribution of the likelihood ratio is $\chi^2$-distributed

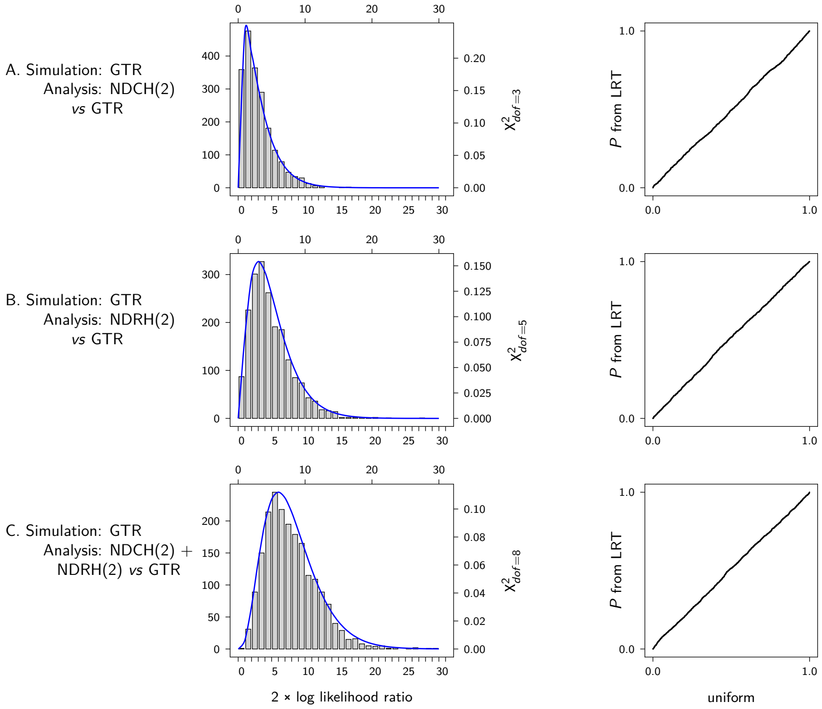


Figure S1: The null distribution of the likelihood ratio is $\chi^2$-distributed when evaluating NDXH models. Simulations (2000 replicates) were made on a 4-taxon trees under the GTR model, and analysed with ML under the GTR and NDXH(2) models as indicated. For the latter, the node for the fourth taxon was assigned one composition vector or exchange rate matrix, and the rest of the tree was assigned the other. Twice the observed log likelihood ratios are shown in the bar plots. The $\chi^2$ PDF is shown as the blue line. Degrees of freedom is 3 for NDCH(2) *vs* GTR, 5 for NDRH(2) *vs* GTR, and 8 for NDCH(2)+NDRH(2) *vs* GTR. PP-plots on the right show $P$-values from the corresponding log likelihood ratio *vs* a uniform distribution, showing that the $\chi^2$ approximation is uniform to a good approximation.

#### LPML effects due to overparameterization using 40-taxon tree simulations

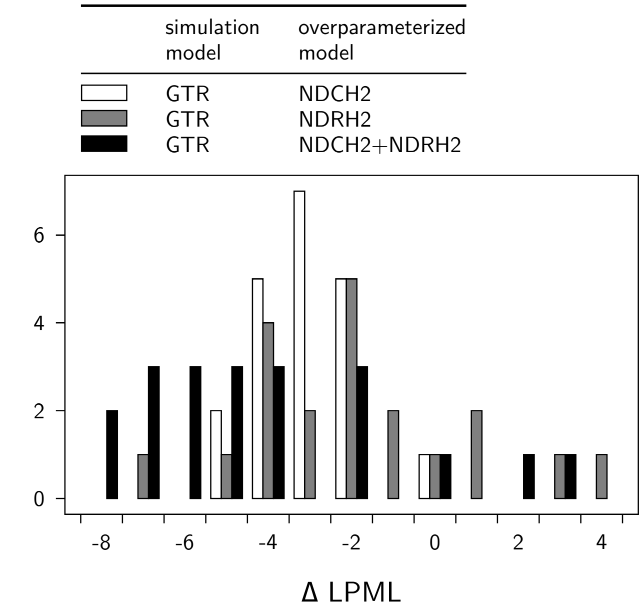


Figure S2: Change in LPML due to overparameterization using data from 40-taxon simulations; similar to Figure 3. MCMC analyses were done with the simulation model and with the overparameterized model. The LPML was calculated for both, and the difference between the overparameterized and the simulation model LPML was plotted. Each condition used 20 replicates. Most differences showed an LPML penalty for overparameterization.

### Cross-estimation between CTH and ERTH

Table S1: LPML values as plotted in Figure 6

| simulation | analysis | LPML |
|---|---|---|
| A: no CTH, noERTH | GTR | -781778.9 |
| | NDCH2 | -781781.2 |
| | NDRH2 | -781784.9 |
| | NDCH2+NDRH2 | -781787.9 |
| B with CTH, no ERTH | GTR | -863852.4 |
| | NDCH2 | -846562.9 |
| | NDRH2 | -862579.2 |
| | NDCH2+NDRH2 | -846569.2 |
| C no CTH, with ERTH | GTR | -803316.5 |
| | NDCH2 | -802863.0 |
| | NDRH2 | -799442.8 |
| | NDCH2+NDRH2 | -799446.2 |
| D with CTH, with ERTH | GTR | -870039.9 |
| | NDCH2 | -847609.0 |
| | NDRH2 | -863403.4 |
| | NDCH2+NDRH2 | -844462.2 |

### Composition and exchange rate effects in simulations analysed with tree-homogeneous models: LPML

Table S2: Model fit for effects demonstration in Figure 7. LPML values are shown. Datasets for rows B and F were 20000 in length, and 100000 for the other datasets.

| row | homogeneous | heterogeneous | difference |
|---|---|---|---|
| A | -528544.6 | -528552.2 | -7.7 |
| B | -105653.9 | -104559.2 | 1094.8 |
| C | -526947.3 | -522177.2 | 4770.2 |
| D | -533704.7 | -515525.2 | 18179.5 |
| E | -534398.5 | -516616.4 | 17782.1 |
| F | -105268.5 | -105141.3 | 127.2 |
| G | -509403.3 | -505235.3 | 4167.9 |
| H | -525324.7 | -518963.7 | 6360.9 |
| I | -514203.0 | -489709.4 | 24493.6 |

#### Composition and exchange rate effects in simulations analysed with tree-homogeneous models: posterior predictive simulations

Table S3: Model fit of CTH as shown in the effects demonstration in Figure 7. Posterior predictive simulations used $X^2$ as the test quantity to show compositional heterogeneity over the tree. Tail area probabilities are shown for the tree-homogeneous model and the tree-heterogeneous model. Row A is from a homogeneous control simulation, and lack of fit is neither expected nor observed. Rows B–E used CTH in their simulations, and the tree-homogeneous model does not fit (P=0). The tree-heterogeneous model fits by this measure. Rows F – I are from simulations that had ERTH and not CTH, and lack of fit of the CTH aspect of the model was not seen in these demonstrations.

| row | homogeneous | heterogeneous |
|---|---:|---:|
| A | 0.995 | 1.000 |
| B | 0.000 | 0.513 |
| C | 0.000 | 0.505 |
| D | 0.000 | 0.404 |
| E | 0.000 | 0.471 |
| F | 0.965 | 0.997 |
| G | 0.293 | 0.939 |
| H | 0.488 | 0.945 |
| I | 0.718 | 0.978 |

#### Cox et al RRNA

Table S4: LPML values for Cox et al analysis. The first set of four values are as shown in Figure 12. These are followed by four sets of LPML values from controls.

| Dataset | analysis model | LPML |
|---|---|---|
| original | GTR | -24035.7 |
| RRNA | NDCH2 | -23627.6 |
| | NDRH2 | -23953.6 |
| | NDCH2 + NDRH2 | -23564.5 |
| control: | GTR | -25359.6 |
| GTR | NDCH2 | -25360.1 |
| no CTH | NDRH2 | -25359.8 |
| no ERTH | NDCH2 + NDRH2 | -25359.0 |
| control: | GTR | -26287.5 |
| with CTH | NDCH2 | -26070.3 |
| no ERTH | NDRH2 | -26281.8 |
| | NDCH2 + NDRH2 | -26072.9 |
| control: | GTR | -25808.1 |
| no CTH | NDCH2 | -25796.9 |
| with ERTH | NDRH2 | -25584.6 |
| | NDCH2 + NDRH2 | -25587.1 |
| control: | GTR | -26817.7 |
| with CTH | NDCH2 | -26625.1 |
| with ERTH | NDRH2 | -26600.3 |
| | NDCH2 + NDRH2 | -26453.4 |

Table S5: Posterior predictive simulations of Cox et al Tree of Life rRNA analyses. The test quantity was $X^2$, and tail area probability is shown. These results show that the analyses using the NDCH2 model fit the composition of the data, but the GTR and NDRH2-only models did not because they do not accommodate CTH.

| | SSU | LSU |
|---|---|---|
| GTR | 0.000 | 0.000 |
| NDCH2 | 0.204 | 0.182 |
| NDRH2 | 0.000 | 0.000 |
| NDCH2 + NDRH2 | 0.222 | 0.218 |

#### Brown et al 2012 ND2 analysis

Table S6: Brown et al 2012 ND2 analysis. The first set of four values, from the original dataset, were plotted in Figure 13. Controls the same size as the original data are included.

| Datset | analysis model | LPML |
|---|---|---|
| original | GTR | -24088.2 |
| ND2 | NDCH2 | -23845.0 |
| | NDRH2 | -24049.9 |
| | NDCH2+NDRH2 | -23821.7 |
| control: | GTR | -22905.8 |
| GTR | NDCH2 | -22906.1 |
| no CTH | NDRH2 | -22905.6 |
| no ERTH | NDCH2+NDRH2 | -22906.5 |
| control | GTR | -25186.3 |
| with CTH | NDCH2 | -24675.5 |
| no ERTH | NDRH2 | -25183.1 |
| | NDCH2+NDRH2 | -24677.9 |
| control | GTR | -24938.8 |
| no CTH | NDCH2 | -24929.2 |
| with ERTH | NDRH2 | -24713.8 |
| | NDCH2+NDRH2 | -24715.4 |
| control | GTR | -23475.6 |
| with CTH | NDCH2 | -23161.6 |
| with ERTH | NDRH2 | -23289.8 |
| | NDCH2+NDRH2 | -23000.3 |

Table S7: Posterior predictive simulations for fit of composition for the model for Brown et al 2012 ND2 analysis. Tail area probability (TAP) is shown for posterior predictive simulations using $X^2$ as the test quantity. The GTR and NDRH2 models are significant (P=0) because those models do not accommodate CTH.

| analysis model | TAP |
|---|---|
| GTR | 0.000 |
| NDCH2 | 0.180 |
| NDRH2 | 0.000 |
| NDCH2 + NDRH2 | 0.162 |
